# Genome-wide protein-protein interaction analysis between aquaporins and harpin (HrpZ2) through molecular docking and MD simulations to unravel its role in growth and stress management in tomato

**DOI:** 10.64898/2026.05.04.722745

**Authors:** Kishori Lal, Tushar Sinha, Sahil Anand, Gaurav Kumar, Aditya Mishra, Debashish Dey

**Affiliations:** Laboratory of Plant Biotechnology, School of Biotechnology, Banaras Hindu University, Varanasi, Uttar Pradesh, 221005

**Keywords:** Harpin (HrpZ2), Aquaporins, Protein-protein docking, Molecular dynamic simulation, Binding affinity

## Abstract

HrpZ2, a harpin protein produced by *Pseudomonas syringae*, a gram-negative plant pathogenic bacterium, elicits hypersensitive response and pathogen defense in non-host plants. Harpins from various bacterial sources elicit varying responses in different non-host plants, due to its structural variations, their precise mechanisms of action are not yet completely understood. As per previous reports, harpins from diverse bacterial sources interact with distinctive members of integral membrane proteins, known as aquaporins. For example, harpin (Hpa1*_Xoo_*) interacts with OsPIP1;3 in rice, whereas, in *Arabidopsis* the harpins Hpa1 and HrpZ interacts with AtPIP1;4 and AtPIP1;3 respectively. Here, we conducted the first genome-wide computational screening of protein-protein interactions between HrpZ2 and all 47 members of tomato aquaporins. Molecular docking identified nine interactors across five subfamilies of aquaporins, with HrpZ2 N-terminal residues mediating these interactions. We validated these via molecular dynamics (MD) simulations, principal component analysis, and free energy landscape analysis, assessing the stability (RMSD, RMSF, radius of gyration), dynamics, and affinity (MM-GMSA). PIP complexes, especially PIP2;1 (-460.46 kcal/mol) and PIP1;7 (-303.82 kcal/mol), exhibited superior stability, compactness, and defined energy minima, confirming PIPs as primary sensors of harpins. Non-PIP aquaporins like TIP1;1 and NIP4;1 showed moderate stability, outperforming weaker interactors (SIP2;1, XIP1;5, XIP1;3). These findings provide robust evidence that HrpZ2 preferentially targets PIPs in tomato, while engaging TIPs and NIPs as auxiliary partners. This multifaceted interaction profile of harpins suggests complex plant-pathogen recognition, modulating aquaporin-mediated cellular responses like growth and stress management in plants.

## Introduction

In nature, plants continuously get challenged by a plethora of pathogens during their growth and development. Plant-pathogen arms race is a sophisticated biology where plants defend themselves by using different layers of defense systems. Generally, plants recognize the pathogens by their conserved slowly evolving microbial signatures called pathogen- or microbe-associated molecular patterns (PAMPs or MAMPs) (Anderson et al., 2010). To detect these molecular patterns, plants have receptors called pattern recognition receptors (PRRs) on their cell surfaces. After recognition, plants mount the first line of defense responses, called PAMP-triggered immunity (PTI), such as callose deposition or cell wall modification and production of reactive oxygen species. Successful pathogens by-pass PTI defense system and are able to cause disease in plants. To hijack the host metabolism and suppress its immune system, successful pathogens inject specialized effector protein into the plant cells, which allows pathogens to colonize the plant system, called effector trigger susceptibility (ETS). In this zig-zag model of co-evolution, plants also parallelly got evolved to counter ETS (Anderson et al., 2010; Kaur et al., 2022). Specifically, to recognize these pathogen’s effectors, plants developed intracellular receptors which recognize these effector proteins and mount a rapid, and robust defense response in the form of programmed cell death of highly localized infected cells and/or its surrounding cells, to restrict the spread of the pathogen infection, called as hypersensitive response (HR) (BalintLKurti P, 2019). Gram negative phytopathogens use their sophisticated system, called type III secretion system (T3SS) to deliver these specialized effector molecules like harpins to plant cells, to counter with host defense response and successfully colonize inside the host (Büttner and He, 2009).

Harpins are a group of effector proteins secreted by numerous gram-negative plant-pathogenic bacteria including several species of genus like *Pseudomonas, Xanthomonas, Ralstonia* etc. During plant-pathogen interactions, phytopathogens secrete harpins through T3SS and deliver directly into the plant cytoplasm (He et al, 1993; Jia and Zhu, 2025). The distinctive physiochemical properties of harpins including very few or no cysteine, glycine-rich and heat-stable protein make them unique from other effectors (He et al., 1993; Wei et al., 1992; Wang et al., 2007; Liu et al., 2018). Different harpins from different phytopathogens have been reported, such as HrpN and HrpW from *Erwinia* spp (Barny M-A, 1995; Gaudriault et al., 1998; Kim and Beer, 1998), Hpa1 and HpaG from *Xanthomonas* spp (Li et al., 2015; Chen et al., 2018), PopA1 and PopW from *Ralstonia* spp. (Arlat et al., 1994; Li et al., 2010), HrpZ and HrpZ2 from *Pseudomonas* spp. (He et al., 1993; Lal et al, 2026). In 2013, Choi et al., classified harpins into four groups based on their functional characterization such as sequence similarity and domain architecture (Choi et al., 2013).

Aquaporins (AQPs, popularly known as water channels) belong to major intrinsic protein (MIP) superfamily expressed almost every living organism majorly plants and animals (Kourghi et al., 2018). In animals, AQPs are crucial for maintaining fluid balance and supporting specialized physiological processes like water reabsorption in kidney, fluid secretion and organ homeostasis (eye, brain, skin etc) (Verkman A. S, 2008). In plants, it shows a large diversity compared to animals. It plays crucial roles in plants like from seed germination, cell elongation, flowering, management of water to nutrient uptake (Wang et al., 2020). Several aquaporin members such as PIPs and TIPs play essential roles in plant growth and development such as OsPIP1;1 promotes seed germination and provide salt resistance in rice (Liu et al., 2013). PIPs and TIPs are also involved in the root growth as they showed high activities in growing root zones in maize (Hukin et al., 2002, Barrieu et al., 1998). Additionally, the aquaporins also improves crop yield by increasing the mesophyll CO_2_ conductance along with essential roles in transporting of phloem sucrose (Xu et al., 2019). Besides, it also plays crucial roles in plant-pathogen interaction via interaction with bacterial elicitors, such as harpin, thereby promoting plant growth and development. It is also involved in stress management by inducing the innate immunity in the non-host plants (Tian et al., 2016; Afzal et al., 2016).

Harpin from different phytopathogens are found to interact with the same or different plant AQPs, may be due to their structural variability and the orientation of interaction interfaces. Harpin such as Hpa1 protein is reported to interact with members of PIP subfamily of aquaporins, such as it interacts with PIP1;4 in *Arabidopsis* (Li et al., 2015) while in Rice it interacts with PIP1; 3 (Li et al., 2019). However, in case of other harpins, such as HrpZ from *P. syringae* pv. *syringae*, it shows an interaction with PIP1;3 in *Arabidopsis* (Lal et al., 2023). Hence, different harpins so far have been reported to interact with only members of PIP subfamily of aquaporins in different non-host plants. The downstream signalling mechanism behind the harpin action, by which they activate plant defences, is not completely characterised. The use of docking algorithms to model protein–protein interactions (PPIs) is cost-effective computational method for understanding biomolecular interactions and mechanisms. By simulating the physical and chemical complementarity between surfaces, these tools can pinpoint specific hot spots, critical residues that drive binding affinity. Molecular docking helps to identify the best native-like protein-protein docking pose from a library of generated decoys requiring a hierarchical approach that combines rapid ranking by scoring functions with computationally rigorous refinement. Further task is to be identifying the important residues present at the protein interfaces, which actively participates to stabilize the protein–protein interaction.

Here, in this study, we conducted the protein-protein molecular docking and molecular simulations analysis to detect possible interaction of harpin (HrpZ2) from *P. syringae* with different members of aquaporins belonging to five different subfamilies of tomato aquaporins, to unravel whether harpins show interaction with other aquaporin members belonging to different subfamilies, apart from PIPs, in tomato. We studied genome-wide 47 members of AQPs in tomato, categorised into five different subfamilies (PIPs – 14 members, TIPs – 11 members, NIPs – 12 members, SIPs – 4 members, XIPs – 6 members). In this investigation, the protein-protein molecular docking and molecular dynamics simulations were performed to study the protein-protein interaction study between harpin (HrpZ2) and AQPs.

## Materials and Methods

### Retrieval of aquaporins sequence from the Sol Genomic and NCBI data bases

A tomato genome-wide searching approach was used for the retrieval of aquaporins (AQPs) sequences from Solanaceae Genomics Network, popularly known as Sol Genomics Network (Solanum pan genomics) (https://solgenomics.net) while the National Centre for Biotechnology Information (NCBI) (https://www.ncbi.nlm.nih.gov/) database was used for the retrieval of amino acid sequences of harpin protein (HrpZ2) from *Pseudomonas syringae* (NCBI gene bank ID: WDY61303.1) respectively, these sequences were used for further studies. Genome-wide searching resulted in a total of 47 members of aquaporin proteins in tomato.

### Prediction of three-dimensional structure, refinement and quality assessment of predicted models of harpin (HrpZ2) and aquaporins

The three-dimensional model of all members of aquaporins from *Solanum lycopersicum* and harpin (HrpZ2) from *P. syringae* were generated using the I-TASSER server (https://aideepmed.com/I-TASSER/) (Zheng et al., 2026). The amino acid sequences were used to predict the 3D-structure of the proteins, in the I-TASSER server. Structural validation and quality assessment of the predicted protein models were done using ERRAT (Colovos and Yeates, 1993), VERIFY 3D (Bowie et al., 1991), and PROCHECK (Laskowski et al., 1996) programs of interactive validation server, SAVES v6.1 (https://saves.mbi.ucla.edu). Finally, the atomic-level refinement of predicted 3D-models were done by using ModRefiner server (https://aideepmed.com/ModRefiner/), where we submitted the 3D-models of proteins in the form of PDB file format as an input file for the refinement process. Meanwhile the Protein Structure Analysis (ProSA), web-based server (https://prosa.services.came.sbg.ac.at/prosa.php) was used to evaluate the predicted 3D structure of proteins based on the predicted Z-score value.

### Physicochemical property analysis

The various physicochemical properties of all the members of aquaporins were estimated using the ProtParam tool of the ExPASy server (https://www.expasy.org) (Gasteiger et al., 2005). To calculate the physicochemical properties, we used the amino acids sequences of the proteins. Various physical and chemical parameters were computed including the amino acid composition, molecular weight, atomic composition, extinction coefficient, aliphatic index, instability index, grand average of hydropathicity (GRAVY) and the estimated half-life.

### Screening of tomato aquaporins for interaction with harpin from *P. syringae* through protein-protein docking

To investigate which aquaporin members are the most possible interactors with or the best binding partner of harpin (HrpZ2), a protein-protein docking was performed with all the members of aquaporins in tomato. The docking was done using the Cluspro 2.0 (https://cluspro.bu.edu/home.php) (Jones et al., 2022), a docking web server. The input files of each protein were uploaded in the form of PDB file format. Cluspro initially uses fast fourier transformation (FFT) based protein-protein interactions program, named as PIPER. In this interaction, former screening was done *via* testing billions of potential conformations of the ligands binding to the receptor. After that, the generated structures were filtered out on the basis of energy. Based on the scoring function, 1000 lowest energy structures were selected. The selected (1000) top-ranked structures were clustered based on their structural similarities (Root Mean Square Deviations). The largest clusters usually represent the most probable near-native biological relevant docking poses. Finally, to fix the minor steric clashes, top-ranked models from the largest clusters were refined as per energy minimization using the CHARMM force field. Docking was performed between harpin (HrpZ2) and one aquaporin protein at a time.

### Interacting residues and computational alanine scanning

A computational approach, alanine scanning was performed using the PPCheck web server (http://caps.ncbs.res.in/ppcheck/) (Sukhwal and Sowdhamini, 2015) to find out key residues which significantly contribute to strengthen the protein–protein docked complex. This approach involves substituting interfacial residues with alanine and calculating the resulting changes after mutation. The total interaction energy of the complex was calculated through total binding energy (kJ/mol) change. It serves as a quantitative measure of each residue’s contribution to the complex stability. Residues that, upon mutation, exhibit a positive change in energy (ΔΔG > 0) indicate that the native residue is important, and it stabilizes the interaction and can therefore be classified as energetically important “hotspot” residue, residues showing a negative or negligible change in energy after mutation suggest that the original residue contributes little to none in the stability of the PPI complex. Overall, alanine scanning with PPCheck provides a robust computational strategy for mapping energetically crucial residues at protein–protein interfaces.

### Stability and binding affinity

After finishing the docking of each member of aquaporins with harpin, stability and binding affinity of each docked complex was evaluated by estimating the free binding energy of the docked complexes to predict the binding affinity of harpin against aquaporin proteins. The Molecular Mechanics with Generalized Born and Surface Area (MM/GBSA) (http://cadd.zju.edu.cn/hawkdock/) (Wang et al., 2021) method was used to calculate the global binding free energy (ΔG_bind (MM-GBSA)_) of the docked complexes. In the MM/GBSA method, the total binding free energy is determined by the following equation:

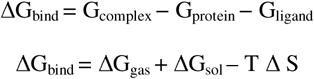

where G_complex_ is corresponds to the free energy of the complexes, whereas G_protein_ and G_ligand_ are represents free energies of the individual partner (protein/ligands). Entropy after ligand binding is represented by TΔS. ΔG_gas_ is corresponds to the total internal energies including angle, bond, dihedral (ΔE_int_), electrostatic component (ΔE_ele_), van der Waals energy (ΔE_vdW_) and ΔG_sol_ (polar component - ΔE_GB_, non-polar component -ΔE_SURF_) of solvation energy.

### Visualization and intramolecular interaction

The harpin-aquaporin docked complexes were selected based on high binding affinity and pictured via BIOVIA Discovery Studio 2021 (https://www.3ds.com/products/biovia/discovery-studio/visualization) and PyMol Version 3.1.0 (open source) (https://pymol.org/2/).

### Molecular Dynamics Simulation of harpin-aquaporins docked complexes

The selected protein-protein docked complexes, were subjected to molecular dynamics (MD) simulation using GROMACS (https://www.gromacs.org) software package through its web-based implementation. The optimized potential for liquid simulations (OPLS) force field pre-processing parameter was used to assess the molecular interaction of the docked complexes while performing the MD simulations. The system was solvated with a simple point charge (SPC) water molecules and placed within a triclinic box with periodic boundary conditions applied in all directions. To ensure electromagnetic neutrality, the system was counterbalanced *via* counter ion (Na^+^ or Cl^-^) to replace corresponding solvent molecules. Initially steepest descent algorithm was applied to remove the unfavourable clashes. In minimization, the system was balanced in two steps: in first step, an thermal equilibration run (where constant number of particles, volume, and temperature of 300 K (NVT)) to stabilize the system temperature, subsequent second equilibration was performed under a constant number of particles, pressure of 1 bar and temperature (NPT) to reach the target density and pressure. Upon successful equilibration, a production MD simulation was carried out for a total duration of 200 ns. Particle mesh Ewald (PME) method was utilized to treat the long-range electrostatic interactions, while 1.0 nm cutoff distance applied to calculates van der waals interactions. The equations of motion were integrated using a standard leap-frog algorithm, and trajectories were saved periodically for subsequent structural and thermodynamic analysis. Finally, the root mean square deviation (RMSD), root mean square fluctuation (RMSF) analysis and Radius of Gyration (Rg) analysis was done.

### Sequence alignments and Phylogenetic analysis of aquaporins proteins

For phylogenetic analysis, the full length amino acid sequences of all the aquaporin (47 members) proteins were retrieved from the Sol genome database and multiple sequence alignment was performed *via* uploading the amino acids sequences in FASTA format and using MEGA 11 (https://www.megasoftware.net) software with clustalW method, followed by building the phylogenetic tree based on Maximum Likelihood method and Jones-Taylor-Thornton model. The tree with the maximum log likelihood with 1000 bootstrap value in the MEGA software was constructed and visualized through interactive tree of life (iTOL) (https://itol.embl.de) (Letunic and Bork, 2024) web server.

### Motif analysis

The conserved motif of aquaporin proteins was identified using the Multiple Expectation Maximization for Motif Elicitation (MEME Suite) (https://meme-suite.org/meme/tools/meme) version 5.5.9 web-based server (Bailey et al., 2015). All the 47 aquaporin proteins were directly submitted in MEME suite server via input sequences file as FASTA format. The motifs were identified by selecting the parameters including the classic mode of motif discovery mode, with the number of motifs required is 10, minimum width of the motifs is 6 and maximum 50.

## Results

### Analysis of predicted three-dimensional structure, refinement and quality assessment of predicted models of harpin (HrpZ2) and aquaporins

I-TASSER is a widely used bioinformatics server for the structure prediction. It predicts 3D-model structures from amino acid sequences following a hierarchical approach, based on iterative threading assembly refinement (I-TASSER). This approach recognizes structural scaffolds from the protein data bank (PDB) using protein treading algorithms. Then constructs full-length 3D-structures by reassembling structural fragments. Finally, these models are refined using atomic level simulations and clustering to generate up to five high quality structural predictions. The quality assessment of the predicted proteins models for harpin and aquaporins (47 members) were done by the hosted tools of SAVES v6 server such as ERRAT, VERIFY 3D, and PROCHECK. The 3D model of harpin and aquaporin members showed that more than 93% of the total residues resided in the allowed and additionally allowed regions while only 0.5% to 6% residues remained in the disallowed region on Ramachandran plots (Fig. 1). The predicted protein models were also assessed for their quality by using ProSA server. The harpin protein showed a Z-score value of -4.88 (Fig. 1A) while the Z-score values for some of the representative members from each subfamily of aquaporins are as -4.39, -4.59, -3.91, -0.95, -1.69, -3.04, -3.8, -2.75, -3.6 for the i) PIP1;7, ii) PIP2;1, iii) TIP1;2, iv) TIP2;2, v) NIP4;1, vi) SIP2;1, vii) XIP1;1, viii) XIP1;3, ix) XIP1;5 protein structures respectively, as shown in Fig. 1.

**Fig. 1.**
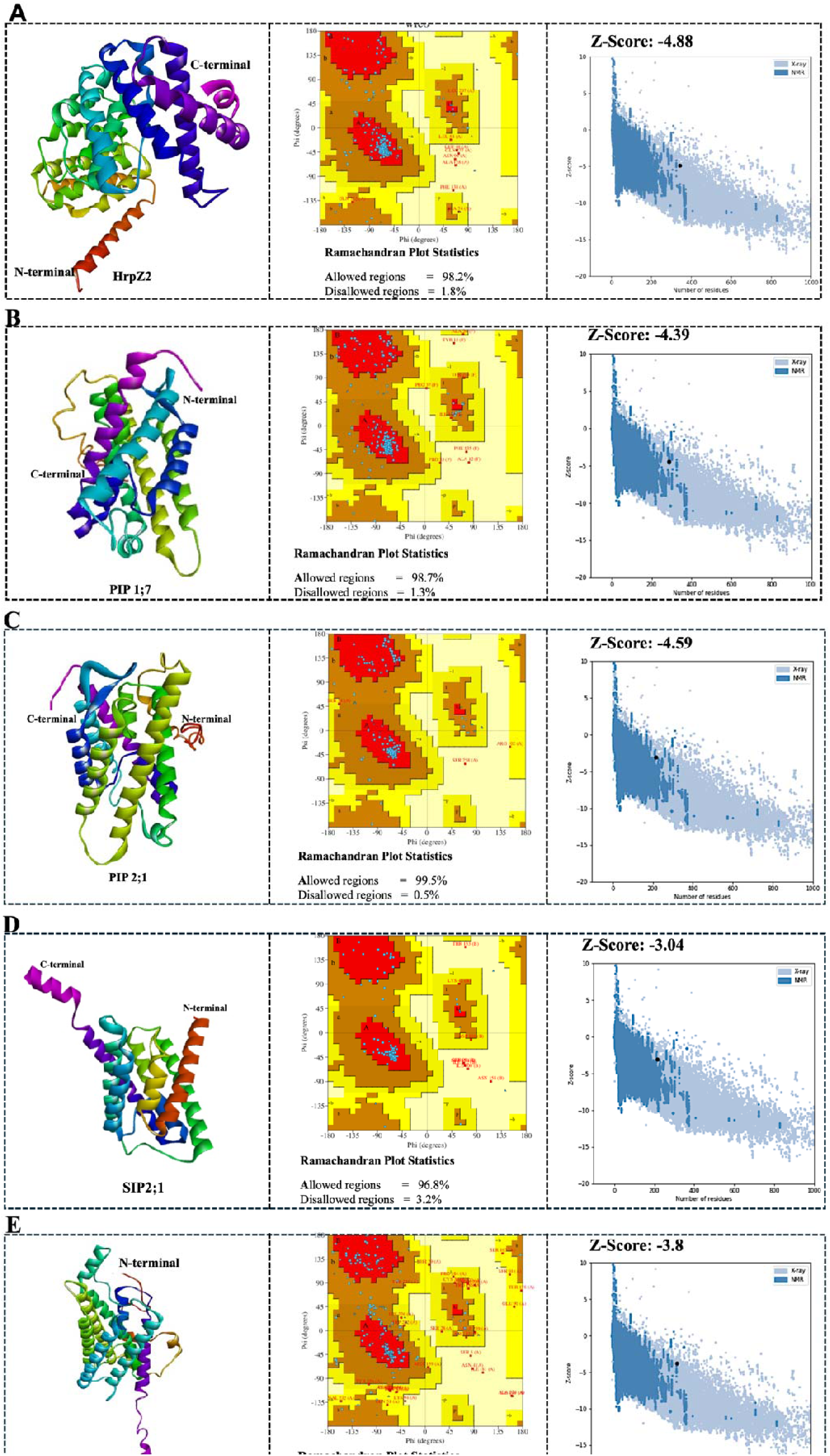
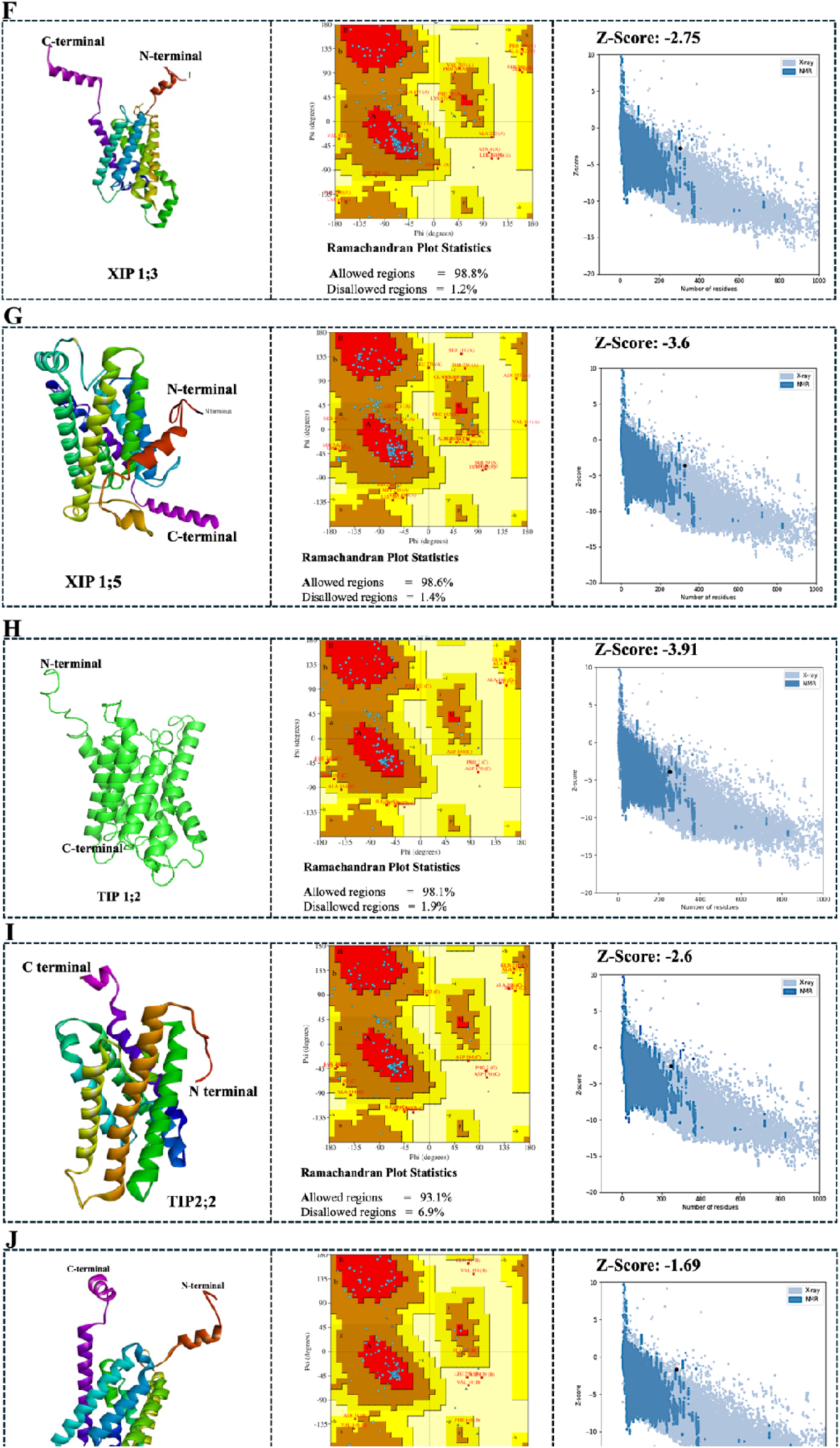
Selected 3D models of harpin (HrpZ2) and aquaporin proteins and their quality assessment. **A.** Predicted 3D model of harpin (HrpZ2) protein followed by model quality by means of Ramachandran plot and Z-score. **B-J.** Predicted 3D models of selected members of aquaporin proteins PIP1;7, PIP2;1, SIP2;1, XIP1;1, XIP1;3, XIP1;5, TIP1;2, TIP2;2 and NIP4;1 followed by analysis of their quality using Ramachandran plot and Z-score.

### Physicochemical property analysis

The various physicochemical properties of aquaporins proteins (47 members) were estimated using the ProtParam tool of the ExPASy server. The aquaporins are small membrane proteins, typically containing 6 membranae-spanning alpha helices. The **Table 1** provides a detailed physicochemical profile of aquaporin proteins in tomato, categorised by their subfamilies including plasma membrane intrinsic proteins (PIPs), tonoplast intrinsic proteins (TIPs), nod-like intrinsic proteins (NIPs), small basic intrinsic proteins (SIPs), and X-intrinsic proteins (XIPs). The analysis showed a significant variation based on their localization. Aquaporins showed a range of molecular weight from 21 to 35 kDa except TIP2;2, (∼136 kDa), NIP4;3, (∼14 kDa), SIP1;3 (∼10 kDa) and XIP1;6 (∼57 kDa). Isoelectric point is one of the important characters of any protein to perform a function. The pI values varied significantly in different aquaporin subfamilies. PIPs, NIPs, and SIPs are generally basic (>7.0), TIPs show a broader range, including several acidic isoforms like TIP1;2 (pI 5.36), TIP4;1 (pI 5.79), and TIP2;3 (pI 5.67) while the XIPs nearly neutral (pI 6-7). The majority of the aquaporins are classified as stable (< 40) except SIP2;1 and XIP1;6 (> 40) which are predicted to be unstable. All the aquaporins members showed positive hydrophobicity (GRAVY) scores confirming their identity as integral membrane proteins.

**Table 1.**
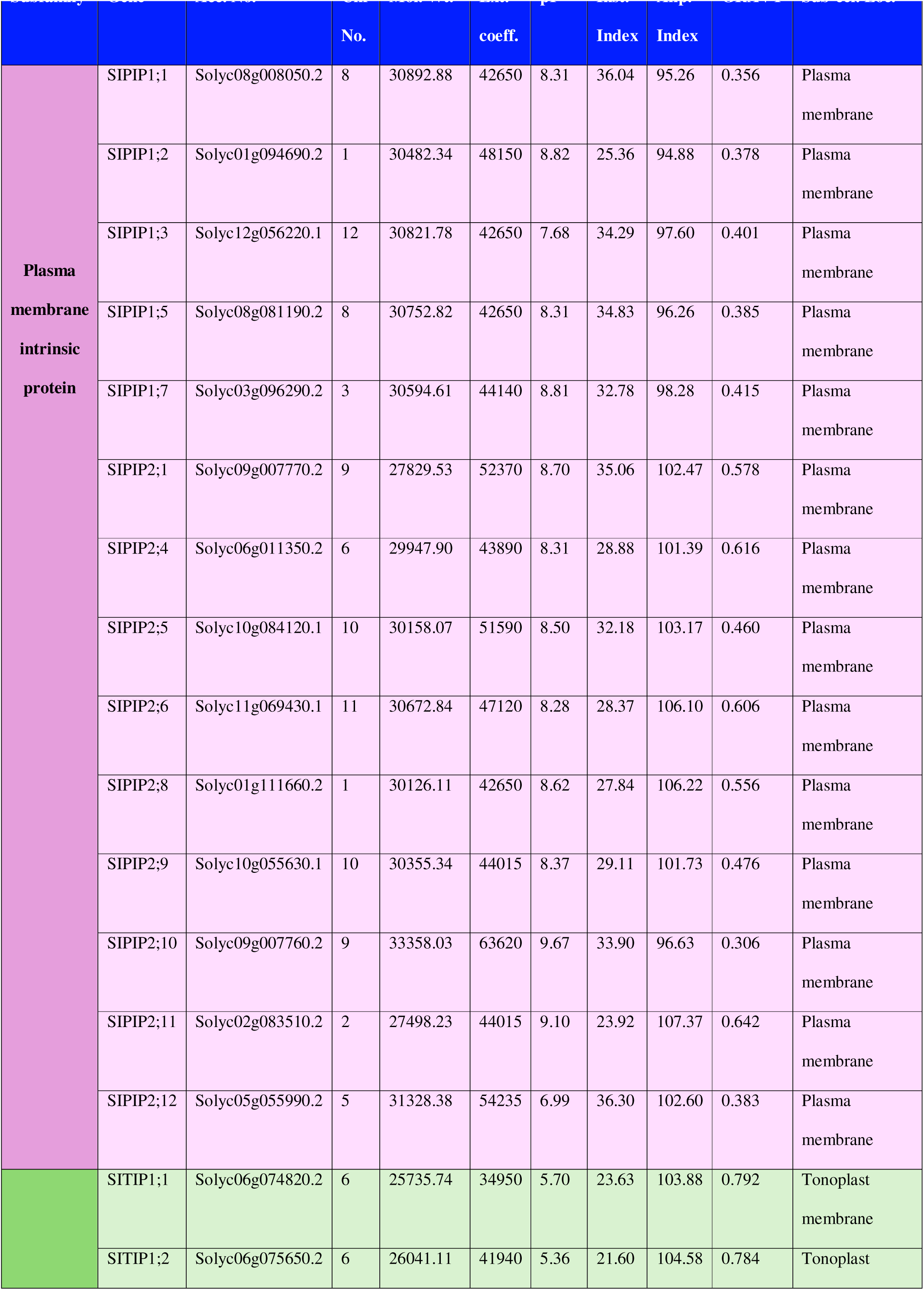
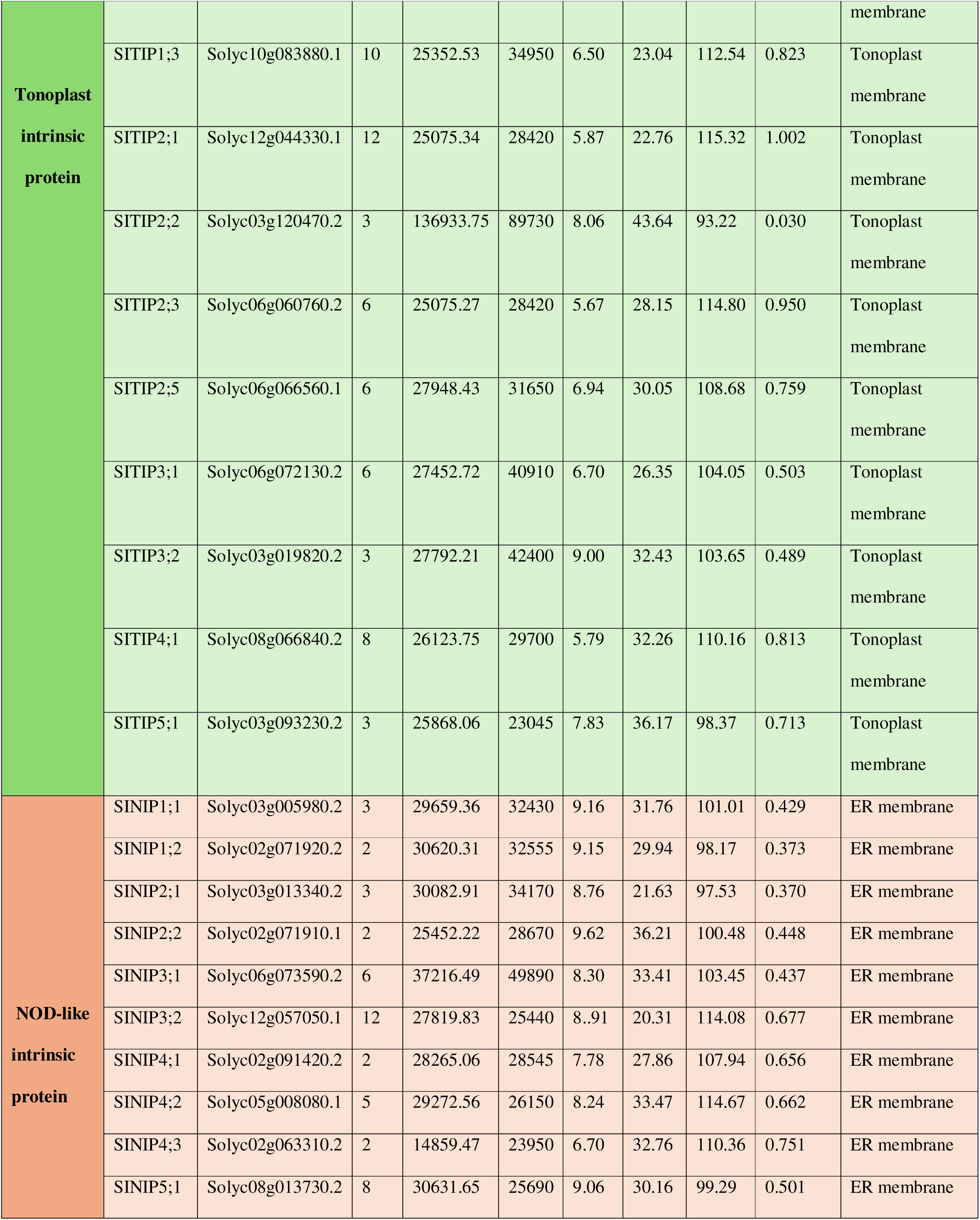
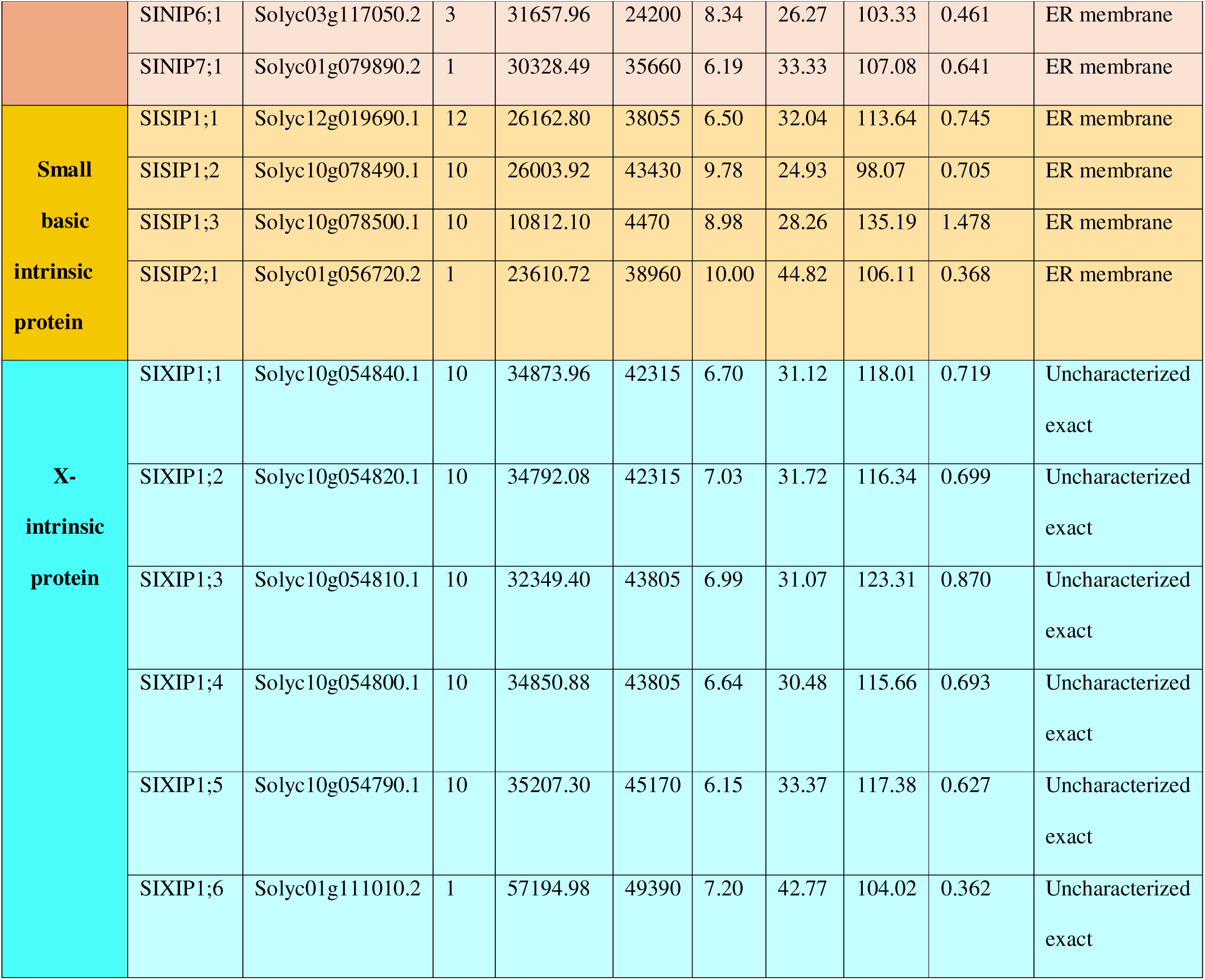
Physicochemical properties of all the aquaporin proteins (47 members) in *Solanum lycopersicum*.

### Analysis of the protein-protein docked complexes of harpin (HrpZ2)-aquaporins and their binding affinity

Genome-wide analysis of interaction of harpin with various members of aquaporins was performed to check whether the harpin binds with the aquaporins of other subfamilies such TIPs, NIPs, SIPs and XIPs, apart from its so far documented interaction only with the members of PIPs subfamily of aquaporins. In this protein-protein docking study, all the 47 members of tomato aquaporins were docked with the predicted model of harpin. Cluspro web server was used to perform all the docking study. After docking, the docked complexes were analyzed and visualized using PyMoLe visualization tool. The genome-wide docking was performed to screen based on their binding free energy score. Initially, the binding free energies of all the docked complexes were predicted *via* the MM/GBSA server. Finally, by comparing the binding free energies of all the aquaporins-harpin docked complexes, we selected only nine top-ranked docked complexes (Table 2) which showed the promising free energy values comparative to other members of aquaporins. The binding free energy values of nine top-ranked docked complexes, as predicted by the MM/GBSA server, are -176.25, -184.57, -460.46, -158.55, -87.44, -198.88, -135.16, - 300.76, and -303.82 for the respective docked complexes of XIP1;5-HrpZ2, SIP2;1-HrpZ2, PIP2;1-HrpZ2, XIP1;1-HrpZ2, TIP2;2-HrpZ2, XIP1;3-HrpZ2, TIP1;1-HrpZ2, NIP4;1-HrpZ2, and PIP17- HrpZ2. Finally, these nine top-ranked docked complexes were selected for further analysis (Table 2).

**Table 2.** Binding free energy of the selected docked complexes predicted by the MM/GBSA server.

| Sr. no. | Selected complexes | Protein–protein complex | MMGBSA $\Delta G$ (kcal/mol) |
| --- | --- | --- | --- |
| 1) | Complex-1 | HrpZ2-PIP2;1 | -460.46 |
| 2) | Complex-2 | HrpZ2-PIP1;7 | -303.82 |
| 3) | Complex-3 | HrpZ2- NIP 4;1 | -300.76 |
| 4) | Complex-4 | HrpZ2-XIP1;3 | -198.88 |
| 5) | Complex-5 | HrpZ2-SIP2;1 | -184.57 |
| 6) | Complex-6 | HrpZ2-XIP1;5 | -176.25 |
| 7) | Complex-7 | HrpZ2-XIP1;1 | -158.55 |
| 8) | Complex-8 | HrpZ2-TIP1;1 | -135.16 |
| 9) | Complex-9 | HrpZ2-TIP2;2 | -87.44 |

The 9 different docked complexes of harpin and aquaporin proteins were analysed to study the protein-protein interaction, mainly the interaction between the two chains involved in the formation of potential hydrogen bonds, salt bridges, disulphide bonds, and non-bonded contacts. In the docked complexes, the interacting residues were highlighted with different colours with respect to the nature of residues. The positively charged residues like histidine, lysine and arginine were coded with blue colour, the negatively charged residues such as aspartic acid and glutamic acid were coded with red colour, the cysteine were coded with yellow colour while the aromatic, aliphatic and neutral residues were coded with violet, grey and green colours respectively.

The interaction analysis of the docked complexes identified a set of potential residues in harpin (chain A) including Glu24, Lys66, Asn103, Asp142,Leu20, Met16, Gln12, Thr13, Ser5, Ser9, Thr183, Phe186, Ala281 and Ala281 that consistently showed interaction with multiple members of aquaporin subfamilies such as Plasma membrane intrinsic protein (PIP1;7 and PIP2;1), Tonoplast intrinsic protein (TIP2;2), X- intrinsic protein (XIP1;1, XIP1;3 and XIP1;5) and Small basic intrinsic protein (SIP2;1) and NOD-like intrinsic protein (NIP4.1). The residues showing high frequency score and interacting with multiple residues of the different partners, are listed in the Table 3. These residues maintain consistent interaction and form an interactive core which might play a crucial role during the interaction of one protein with their target partner. In case of harpin protein, most of the residues which are consistently involved in the interaction are present at the N-terminal. The interaction interface is composed of different interaction types like potential hydrogen bonding, electrostatic interaction and hydrophobic interactions. While some of the residues showed peripheral or transient interaction with minimal contribution to the binding stability and some missing interaction in certain protein which is likely due to the structural variation and different binding orientations.

**Table 3.**
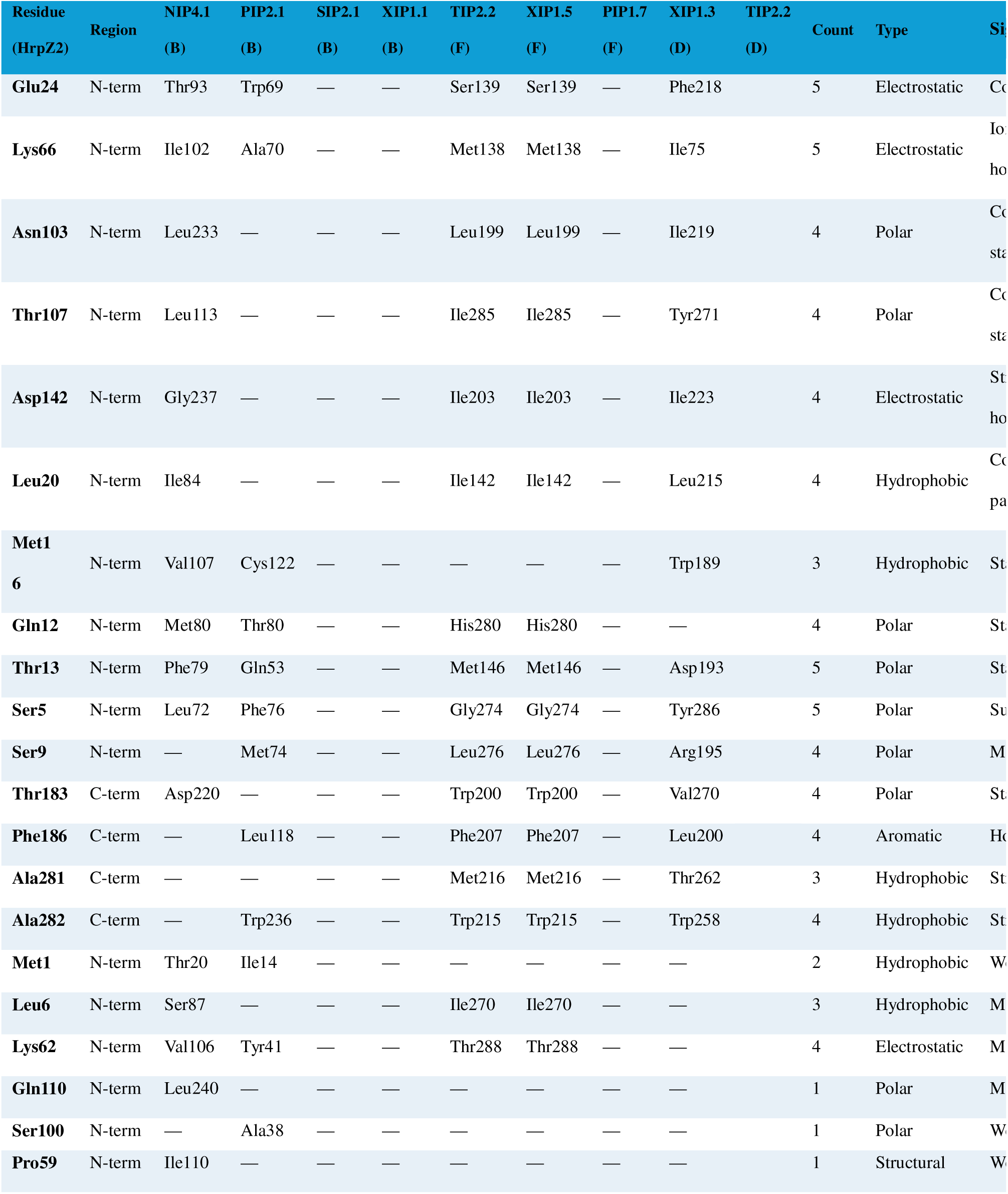
Potential interacting residues and their significance as a hotspot in the harpin-aquaporin interaction study.

The docked complexes of harpin and aquaporins protein were analysed to study the protein-protein interaction. Mainly the interaction between the two chains involved the formation of potential hydrogen bonds, salt bridges, disulphide bonds, and non-bonded contacts. In docked complex, the interacting residues were highlighted with different colours with respect to the nature of residues (Fig. 2). The positively charged residues like histidine, lysine and arginine were coded with blue colour, the negatively charged residues such as aspartic acid and glutamic acid were coded are coded with red colour, the cysteine were coded with yellow colour while the aromatic, aliphatic and neutral residues were coded with violet, grey and green colours respectively. The interaction analysis of the docked complexes identified a set of potential residues in harpin (chain A) including Glu24, Lys66, Asn103, Asp142, Leu20, Met16, Gln12, Thr13, Ser5, Ser9, Thr183, Phe186, Ala281 and Ala281 that consistently interacted with multiple members of aquaporin subfamilies such as Plasma membrane intrinsic protein (PIP1;7 and PIP2;1), Tonoplast intrinsic protein (TIP2;2), X- intrinsic protein (XIP1;1, XIP1;3 and XIP1;5) and Small basic intrinsic protein (SIP2;1) and NOD-like intrinsic protein (NIP4.1). The residues showing high frequency score and interacting with multiple residues of the different partners, are listed in the Table 3. These residues maintain consistent interaction and form an interactive core which might play a crucial role during the interaction of one protein with their target partner. In case of harpin protein the most of the residues which are consistently involved in the interaction are present at the N-terminal. The interaction interface is composed of different interaction types like potential hydrogen bonding, electrostatic interaction and hydrophobic interactions. While some of the residues showed peripheral or transient interaction with minimal contribution to binding stability and some missing interaction in certain protein which is likely due to the structural variation and different binding orientations.

**Fig. 2.**
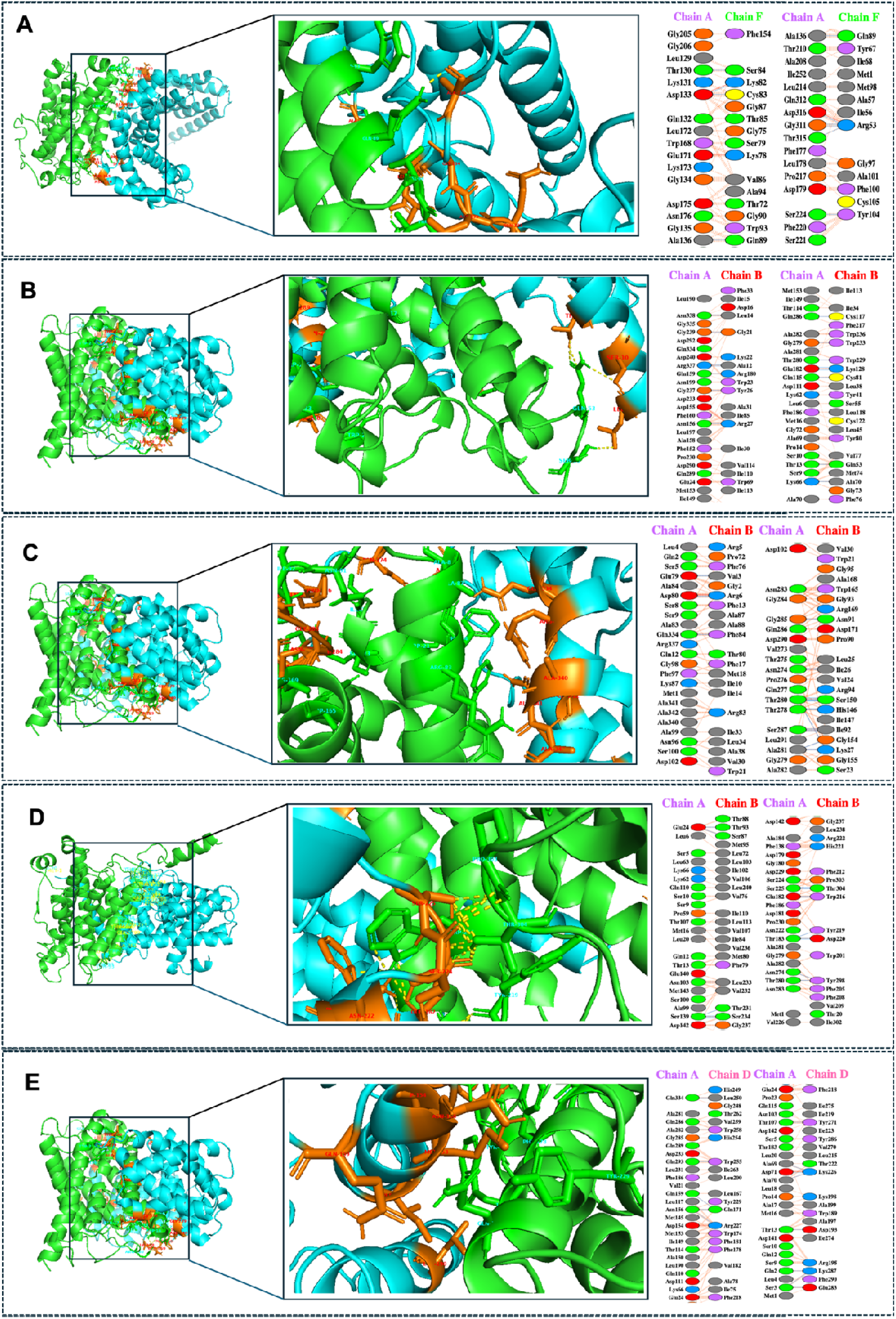
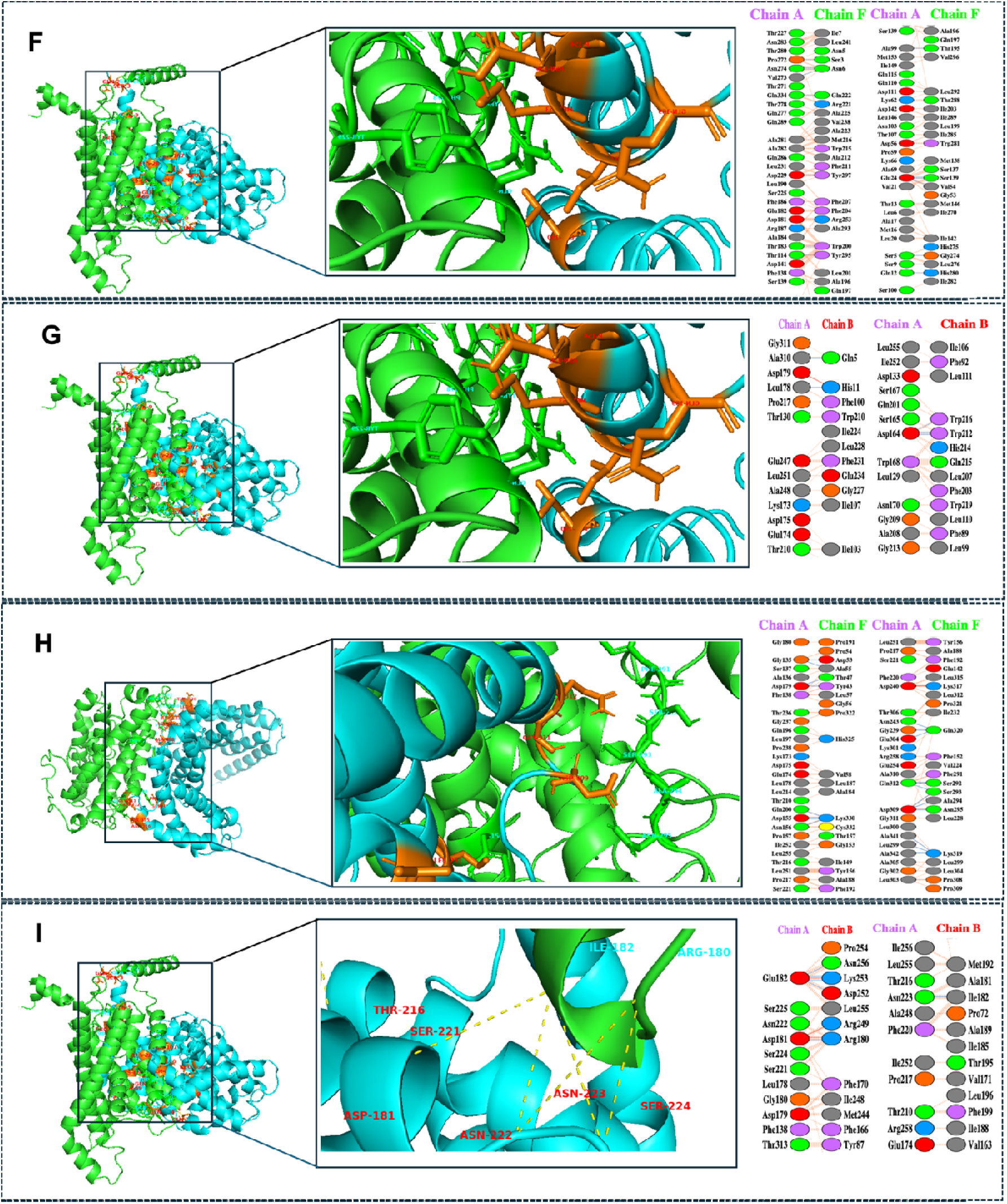
Selected docked complexes (left) of harpin and aquaporin proteins and their interactive residues (Right). **A.** HrpZ2-PIP1;7, **B.** HrpZ2-PIP2;1, **C.** HrpZ2-SIP2;1, **D.** HrpZ2-XIP1;1, **E.** HrpZ2-XIP1;3, **F.** HrpZ2-XIP1;5, **G.** HrpZ2-TIP1;1, **H.** HrpZ2-TIP2;2, and **I.** HrpZ2- NIP 4;1. Docked complexes (left), zoomed view of interaction area in the docked complex (middle) and their interactive residues (Right). In protein-protein interaction the harpin is represented by chain A in all the docked complexes while the other selected aquaporins protein are represented as chain B (NIP4;1, XIP1;1, PIP2;1, SIP2;1), chain D(XIP1;3) and chain F (XIP1;5, PIP1;7, TIP2;2).

### Computational alanine scanning (CAPS) analysis of interacting residues through alanine mutagenesis study

The PPCheck web server simulates the mutation of residues to alanine called alanine scanning mutagenesis to find out how it affects the residue level binding energy of the complex. After the protein-protein docking study the interaction analysis revealed the potential interacting residues involved in the HrpZ2-APQs interaction. Usually PPI contains lots of residues, although only a few residues in the protein-protein interface contribute to the overall stability of binding. Based on the analysis of selected 9 top complexes, we identified a set of potential residues of harpin which contributes to binding (Table 4).

**Table 4.** Computational mutagenesis analysis through alanine-scanning at HrpZ2-aquaporin binding interface.

| Residue<br>(HrpZ2) | PDB-ID | Before Mutation | After Mutation | Binding energy change<br>($\Delta\Delta G_{bind}$) | | Difference |
| --- | --- | --- | --- | --- | --- | --- |
|  |  |  |  | Before<br>Mutation | After<br>Mutation |  |
| **Glu24** | Xip1.5 | A24GLU | A24ALA | -482.45 | -448.93 | 33.52 |
| **Thr183** | Xip1.5 | A183THR | A183ALA | -482.45 | -471.56 | 10.89 |
| **Phe186** | Xip1.5 | A186PHE | A186ALA | -482.45 | -472.31 | 10.14 |
| **Asp142** | Xip1.5 | A142ASP | A142ALA | -482.45 | -477.34 | 5.11 |
| **Gln110** | Xip1.5 | A110GLN | A110ALA | -482.45 | -477.39 | 5.06 |
| **Leu20** | Xip1.5 | A20LEU | A20ALA | -482.45 | -477.84 | 4.61 |
| **Asn103** | Xip1.5 | A103ASN | A103ALA | -482.45 | -478.44 | 4.01 |
| **Gln12** | Xip1.5 | A12GLN | A12ALA | -482.45 | -478.79 | 3.66 |
| **Thr107** | Xip1.5 | A107THR | A107ALA | -482.45 | -478.99 | 3.46 |
| **Lys62** | Xip1.5 | A62LYS | A62ALA | -482.45 | -479.14 | 3.31 |
| **Thr13** | Xip1.5 | A13THR | A13ALA | -482.45 | -479.60 | 2.85 |
| **Ser5** | Xip1.5 | A5SER | A5ALA | -496.75 | -494.44 | 2.31 |
| **Lys66** | Xip1.5 | A66LYS | A66ALA | -482.45 | -480.64 | 1.81 |
| **Met16** | Xip1.5 | A16MET | A16ALA | -482.45 | -480.69 | 1.76 |
| **Pro59** | Xip1.5 | A59PRO | A59ALA | -482.45 | -480.83 | 1.62 |
| **Ser9** | Xip1.5 | A9SER | A9ALA | -482.45 | -480.92 | 1.53 |
| **Met1** | Xip1.5 | A1MET | A1ALA | -482.45 | -481.58 | 0.87 |
| **Ser100** | Xip1.5 | A100SER | A100ALA | -482.45 | -482.11 | 0.34 |
| **Leu6** | Xip1.5 | A6LEU | A6ALA | -496.75 | -496.75 | 0.00 |

The energy contribution of one residue at a time was measured upon mutation to alanine. The interacting residues of harpin at the N-terminal majorly contributes in the interaction and strengthen the harpin-aquaporin complex. The potential residues at the PPI interface such as Glu24, Thr183, Phe186, Asp142, Gln110, Leu20, Asn103, Gln12, Thr107, Lys62, Thr13 and Ser5 contribute a significant change in energy (ΔΔGbind) 33.52, 10.89, 10.14, 5.11, 5.06, 4.61, 4.01, 3.6, 3.46, 3.31, 2.85 and 2.31 respectably.

After alanine scanning, the binding free energy got reduced (less negative) compared to the unmuted (wild) which indicates that mutation destabilized the protein-protein interaction. This suggests that mutated residues contributed favourably to the binding interface, and its substitution with alanine weakened the interaction, resulting in lower binding affinity. Free energy contribution of HrpZ2’s residues in the protein-protein interaction are listed in Table 4.

### Molecular Dynamics Simulation of harpin aquaporins docked complexes

Molecular dynamics simulations are the state of art computational methods used to simulate the time-dependent physical movements of atoms and molecules in biomolecular systems such as the moment and trajectory analysis of the molecules over time. It helps to understand the acceleration (a) on every atom by calculating the force (F) on each atom and update its position and velocity at discrete time steps to capture the fastest molecular vibrations to understand the stability and flexibility of protein-protein docked complexes. In this study, the MD simulations of nine different docked complexes (Table 2) were performed to clarify the stability of the protein-protein docked complexes and flexibility of the protein structures. All the MD simulations were accomplished for 200 ns employing the MD simulation tool unified within SiBIOLEAD (https://sibiolead.com/), which utilizes the GROMACS simulation software.

The structural stability of all the nine complexes was evaluated using RMSD, RMSF, radius of gyration (Rg), solvent-accessible surface area (SASA) and total energy profiles obtained from MD simulation (Fig. 3). RMSD analysis revealed that most complexes reached the equilibrium within the first 20-50 ns, followed by relatively stable trajectories, indicating successful system equilibration and absence of large-scale structural disruption.

**Fig. 3.**
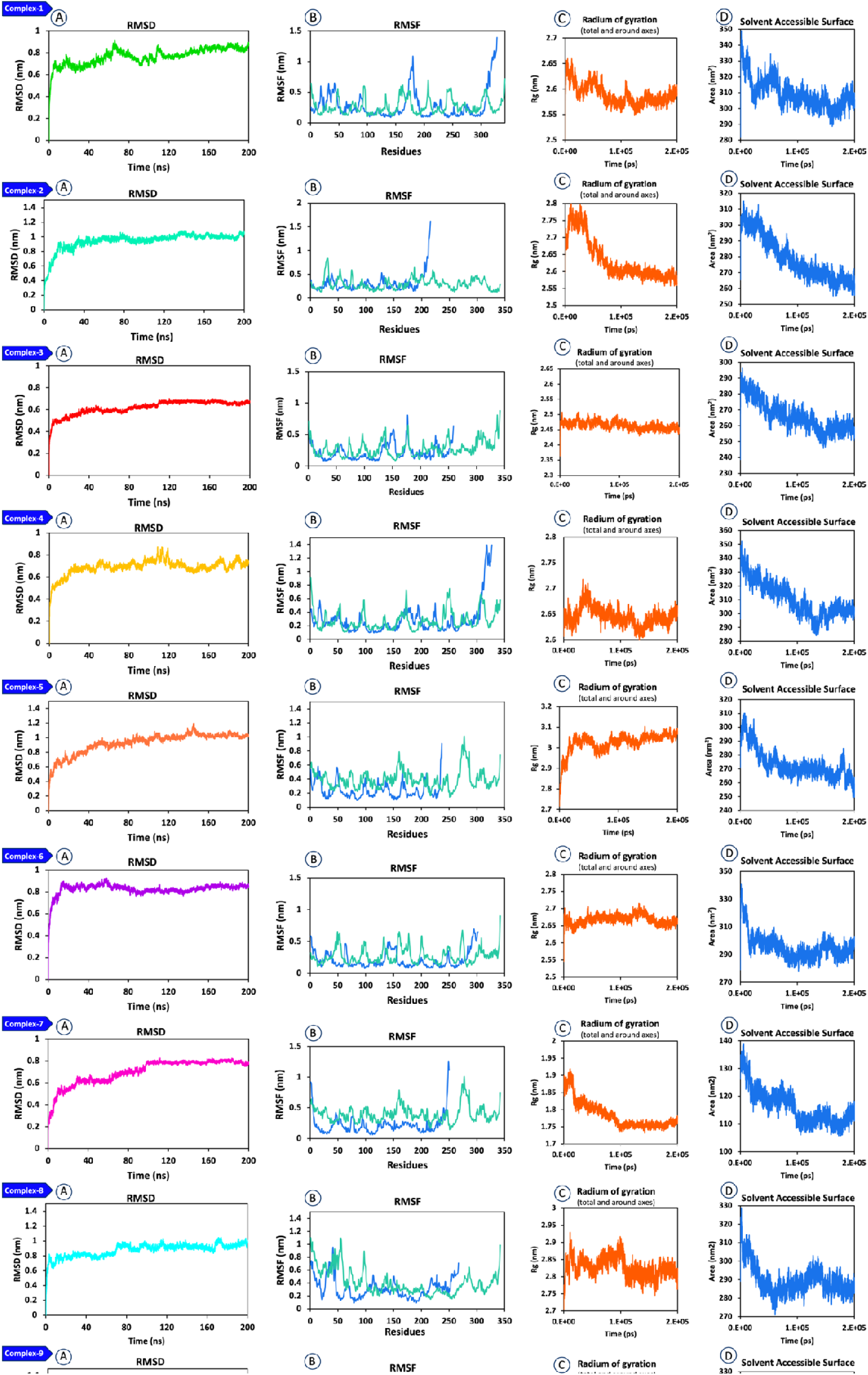
Time dependent molecular dynamics simulation of selected 9 complexes of harpin (HrpZ2) with aquaporins**. A.** Root mean square deviation (RMSD), **B.** Root mean square fluctuation (RMSF), **C.** Gyration (Rg) and **D.** Solvent accessible surface area of complex-1, 2, 3, 4, 5, 6, 7, 8, and 9.

### Root mean square deviation (RMSD) analysis

All complexes achieved equilibrium during the MD simulation; however, their stability profiles differed significantly. Complex-1 (HrpZ2-PIP2;1) and complex-2 (HrpZ2-PIP1;7) displayed stable RMSD trajectories with minimal fluctuations, indicating strong structural integrity. Complex-8 (HrpZ2-TIP1;1) and complex-3 (HrpZ2-NIP4;1) also showed relatively stable behavior. In contrast, complexes involving TIP, XIP and SIP members (Complex-4, 5, 6, 7, 9) exhibited higher RMSD fluctuations, suggesting reduced stability.

### RMSF analysis

RMSF analysis showed that fluctuations were primarily localized to loop regions. Importantly, binding site residues in complex-1 (HrpZ2-PIP2;1) and complex-2 (HrpZ2-PIP1;7) exhibited reduced flexibility, indicating stable ligand engagement. Higher fluctuations were observed in weaker complexes, particularly HrpZ2-SIP2;1 (complex-5) and HrpZ2-XIP1;5 (complex-6).

### Compactness and solvent exposure assessment through radius of Gyration (Rg) and solvent accessible surface area (SASA) analysis

Complex-1 and complex-2 maintained consistent radius of gyration and stable SASA profiles through the simulation, indicating compact and well-folded structures. Complex-8 (HrpZ2-TIP1;1) complex-3 (HrpZ2-NIP4;1) also showed moderate compactness. In contrast, XIP and SIP complexes displayed greater fluctuations, reflecting reduced structural integrity.

### Free energy landscape, principal component analysis and hydrogen bonding analysis

Additionally, the analysis of free energy landscape (FEL) revealed deep and well-defined global minima, indicating thermodynamically favourable and stable conformational states. Importantly, these dynamic and structural observations strongly correlate with the results of MM/GBSA binding free energy analysis, where HrpZ2-PIP2;1 (-460.46 kcal/mol), and HrpZ2-PIP1;7 (-303.82 kcal/mol) exhibited energetically most favoured candidates among all studied complexes. Principal component analysis (PCA) showed that these complexes occupied a restricted conformational space, reflecting limited large-scale motions and enhanced structural rigidity. Hydrogen bond analysis revealed that HrpZ2-PIP2;1 (complex-1) and HrpZ2-PIP1;7 (complex-2) formed the higher number of stable and persistent hydrogen bonds with the harpin elicitor. HrpZ2-TIP1;1 (complex-8) and HrpZ2-XIP1;5 (complex-6) showed moderate hydrogen bonding, whereas weaker complexes exhibited fewer and transient interactions.

### Phylogenetic analysis of aquaporin proteins

The phylogenetic analysis revealed a clear clustering of the identified sequences into five distinct subfamilies, namely NIP, TIP, PIP, SIP, and XIP (Fig. 4). Each subgroup formed well-supported clades, indicating evolutionary conservation and functional specialization within each family. The PIP subfamily showed tight clustering with shorter branch lengths, suggesting a high degree of sequence conservation, which aligns with their essential role in facilitating transmembrane water transport. In contrast, members of the NIP group exhibited relatively longer branch lengths, reflecting greater sequence divergence and potential functional diversity, particularly in the transport of small solutes and metalloids. The TIP and SIP subfamilies also formed distinct clusters, highlighting their evolutionary separation and specialized intracellular roles, such as vacuolar transport (TIPs) and endoplasmic reticulum-associated functions (SIPs).

**Fig. 4.**
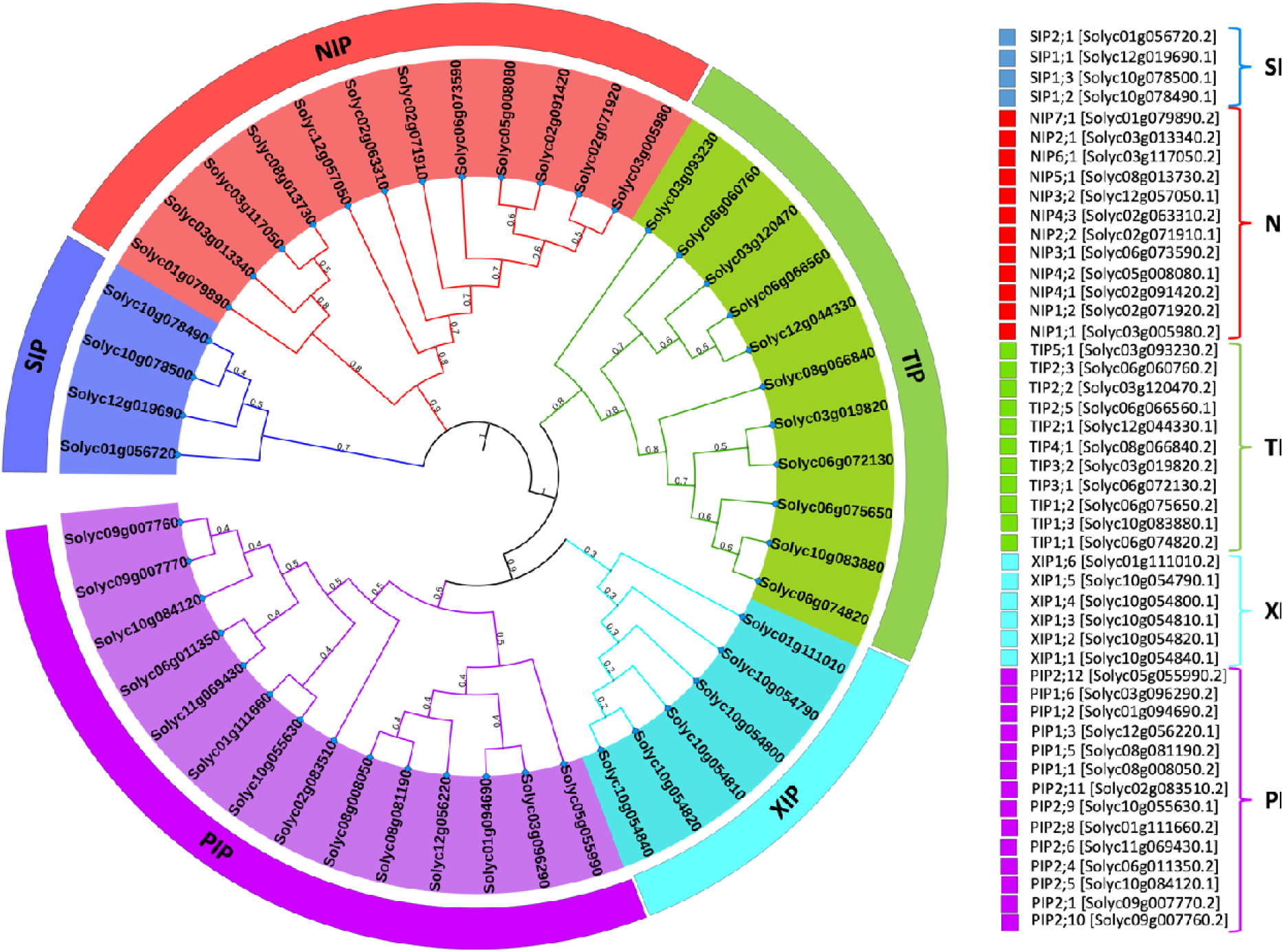
Phylogenetic analysis of all the members of aquaporin proteins from tomato. For the bootstrap test, 1000 replicates were used and next to branches, the branch length is presented. The protein ID and the corresponding accession numbers were displayed at right side along with five subfamilies of aquaporins.

The distribution of aquaporin gene members across different clades suggests possible gene duplication events followed by functional divergence. Overall, the phylogenetic relationships support the hypothesis that aquaporin gene families have evolved through a combination of conservation and diversification to meet diverse physiological demands in plants.

### Conserved motif analysis

The conserved motif analysis revealed a distinct pattern of motif distribution among the identified protein sequences, highlighting both conservation and functional divergence within the gene family. A total of ten conserved motifs were identified (Fig. 6), many of which were widely distributed across most members, indicating their essential role in maintaining the core structural and functional integrity of the proteins. Notably, certain motifs were highly conserved in terms of sequence composition and positional arrangement, suggesting their involvement in key functional domains. Locations and logos of the conserved motif in tomato aquaporins are presented with their statistical e-value, numbers of sites and width of motifs (Fig. 5).

**Fig. 5.**
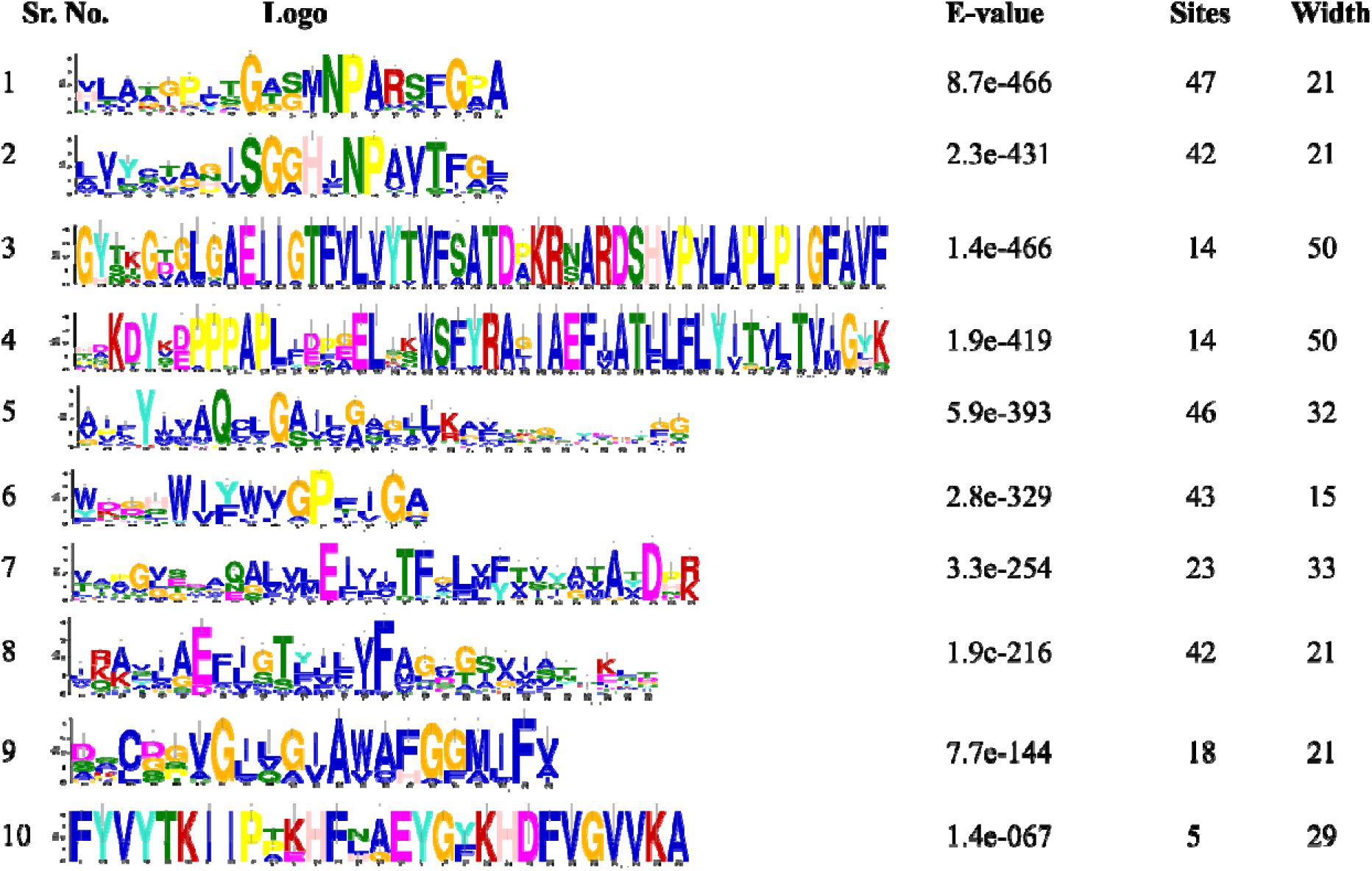
Conserved motif locations in tomato aquaporins as indicated by MEME tools using their amino acid sequences. The 10 sequence logos are presented with their statistical e-value, numbers of sites and width of motifs.

**Fig. 6.**
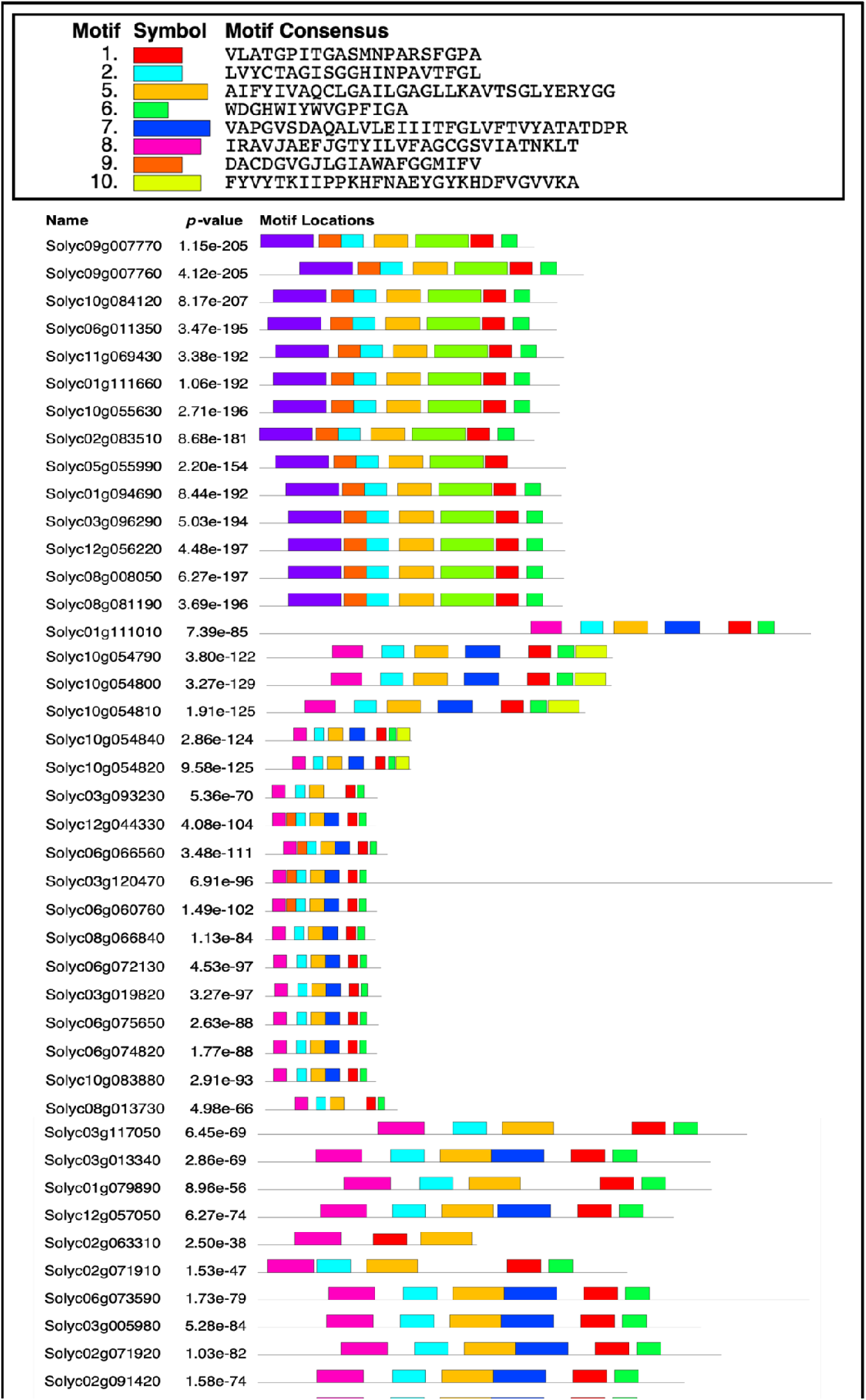
Motif-based sequence analysis of aquaporin proteins. Distribution of the conserved motifs in all 47 aquaporins proteins from tomato. The motifs (10) were color coded (symbol) and amino acid sequences were displayed against each motif. The scale bar at the bottom indicates the protein length.

Members belonging to the same phylogenetic subgroup exhibited similar motif compositions and organization, reinforcing the evolutionary relationships observed in the phylogenetic analysis. For instance, proteins within closely related clades shared common motif architectures, whereas variations in motif sequence (presence, absence, or re-arrangement) were observed across different subfamilies, reflecting functional specialization. Some proteins showed partial or missing motifs, which may indicate sequence divergence, potential functional modification, or gene truncation events. The presence of subgroup-specific motifs further suggests specialized roles of these proteins in different cellular or physiological processes. Additionally, the conserved motifs likely correspond to important structural features such as transmembrane regions or substrate-selective filters, which are critical for the protein activity. Overall, the motif distribution pattern supports the notion that this gene family has evolved through a balance of conservation and diversification, enabling both maintenance of fundamental functions and adaptation to diverse biological roles.

## Discussion

A study of the interaction between a non-host plant (tomato) and pathogen elicitor protein (HrpZ2) is essential to understand the mechanisms of plant defence responses towards bacterial infections. This knowledge may be crucial in the development of disease prevention strategies to fight against phytopathogenic infections and increase the crop yield. HrpZ2 protein is known to induce defence responses in non-host plants, while aquaporin proteins are an integral membrane protein forming pores to transport water and small molecules between the cells. Apart from the role of water regulation in plants, AQPs also perform several other functions in plants, such as cell signalling, nutrient transport, carbon fixation and abiotic and biotic stress responses. The 3D structures of HrpZ2 and 47 AQPs were predicted using I-TASSER, due to unavailability of experimentally derived crystallized structures of HrpZ2 and tomato’s AQPs yet. To overcome this structural strain, various computational tools and algorithms have already been developed which have proven to be an emerging approach to understand the structure-function relation of proteins and their interaction with other protein (Kolodny and Kosloff, 2013; Zhang and Qian, 2024). In this study, PPI between HrpZ2 protein from *P. syringae* and AQPs of *S. lycopersicum* were elucidated using the structural and molecular docking-based approach. At the very beginning, all the predicted 3D structures were subjected to quality assessment with ERRAT, Verify3D, Procheck, tools of SAVES server which showed acceptable Z-score values in the ProSA server (Fig. 1). HrpZ2 structure analysis showed that more than 98% of total residues resided in the allowed region in the Ramachandran plot with a Z-score of (-4.88) while majority of AQPs also showed high quality of structures, with more than 93% total residues residing in the allowed region and acceptable Z-score values. After docking, the Protein-protein docked complexes of HrpZ2-AQPs were evaluated based on their binding free energies. The protein-protein interactions were analysed by MM/GBSA tool to check the binding affinity and stability of the complexes. The complexes which exhibit high binding affinity was selected such as -460.46 kcal/mol, -303.82 kcal/mol, -300.76 kcal/mol, -198.88 kcal/mol, -184.57 kcal/mol, -176.25 kcal/mol, -158.55 kcal/mol, -135.16 kcal/mol and -87.44 kcal/mol displayed by the selected complex-1, complex-1, complex-3, complex-4, complex-5, complex-6, complex-7, complex-8, and complex-9 respectively. Furthermore, molecular dynamics simulations were performed to check the stability of the top ranked protein-protein docked complexes.

In this study, we intended to search for all the possible interactions of harpin (HrpZ2) with different members of tomato aquaporins. We were curious to know whether harpin interacts with members of only one sub-family (PIPs sub-family) of aquaporins, as indicated by various reports till date (Li et al., 2015; Li et al., 2019; Lal et al., 2023) or it may also show possible interaction with members of other subfamily of aquaporins. For the sake of curiosity, we studied *in silico* interaction of harpin with all the 47 members of aquaporin, detected in the tomato genome, which belonged to five different subfamilies of aquaporins (PIPs – 14 members, TIPs – 11 members, NIPs – 12 members, SIPs – 4 members, XIPs – 6 members). In this investigation, the molecular docking and MD simulation were performed to predict the most potential interactor of harpin (HrpZ2) in tomato. The energy contribution of one residue at a time was measured upon mutation to alanine. The interacting residues of harpin at the N-terminal majorly contributes in the binding and stability of the harpin-aquaporin complexes. The potential residues at the PPI interface such as Glu24, Thr183, Phe186, Asp142, Gln110, Leu20, Asn103, Gln12, Thr107, Lys62, Thr13 and Ser5 contributed to the significant change in energy (ΔΔGbind) 33.52, 10.89, 10.14, 5.11, 5.06, 4.61, 4.01, 3.6, 3.46, 3.31, 2.85 and 2.31 respectively. As per the alanine scanning study, the binding free energy got reduced (less negative) as compared to the wild type which indicates that mutation destabilized the protein-protein interaction. This suggests that mutated residues contributed favourably to the binding interface, and its substitution with alanine weakened the interaction, resulting in lower binding affinity. Free energy contribution of HrpZ2’s residues in the protein-protein interaction were listed in Table 4.

The results from the present study provided clear evidence that while the harpin elicitor is capable of binding with multiple aquaporin types, there is a strong preference toward the PIP isoforms. The significantly lower binding free energies observed for PIP2;1 (complex-1) and PIP1;7 (complex-2) indicate that these proteins represented the most thermodynamically acceptable results, where both complexes exhibited stable RMSD profiles, reduced residue-level fluctuation, and well-defined free energy minima. These characteristics are indicative of strong and stable protein-protein interactions. Importantly, the study also demonstrates that non-PIP aquaporins members, particularly TIP1;1 (complex-8) and NIP4;1 (complex-3), may also form moderately stable complexes with the harpin elicitor. This suggests that harpin may interact with different members from multiple aquaporins subfamilies, potentially contributing to a broader-signalling mechanism. However, the comparatively weaker stability and higher conformational variability of XIP and SIP complexes indicate that these are less likely to function as primary interactors for the harpin elicitor. From a mechanistic perspective, the stable binding observed in PIP complexes may facilitate modulation of membrane transport processes, including water and reactive oxygen species (ROS) movement, resulting in tight water regulations in different non-host plants, which are crucial components of plant immune signalling. The involvement of TIP and NIP subfamilies further suggests potential roles of harpin in inducing intracellular signalling pathways.

When the harpin is sprayed on the plant leaves, it is localized in the apoplastic space outside the plasma membrane, where it can be directly sensed only by the aquaporin members of plasma membrane intrinsic membrane (PIPs) subclass, present on the plasma membranes. Accordingly, PIP-type aquaporins are so far reported as the only primary sensors/receptors for extracellular harpin in non-host plants (Li et al., 2015; Lal et al., 2023, Patolia et al., 2024). However, during the plant-pathogen interactions, the harpin is directly secreted into the host plant cytosol by host-specific pathogen through the T3SS system, so only a subset of localized cells may receive harpin in an intracellular context. Once inside the cytosol, harpins may get sensed/perceived by other members of non-PIP aquaporin subclasses, present on the endoplasmic reticulum, tonoplast etc., providing a basis for intracellular sensing of harpins. On this basis, we hypothesize that harpins could be perceived not only in an extracellular form, by the PIP subclass of aquaporins at the plasma membrane but also in an intracellular form by the members of non-PIP aquaporins subfamilies, within the cell. Our molecular docking analysis supports this possibility, as several members of the NIP, XIP, TIP and SIP subfamilies of aquaporins have shown promising interaction with harpin, thereby becoming the harpin-interacting partners, in addition to PIPs.

## Conclusion

This study was designed to identify potential interactors of the bacterial elicitor harpin (HrpZ2) protein in tomato, with a particular focus on the members of aquaporin superfamily. While previous reports suggested that harpins from different bacterial origin, primarily interacts only with the plasma membrane intrinsic proteins (PIPs), the possibility of interactions with other aquaporin subfamilies (TIPs, NIPs, SIPs, and XIPs) remained unexplored. To address this, a comprehensive computational workflow combining molecular docking, binding free energy estimation, and molecular dynamics (MD) simulations was employed. Initial docking and binding energy screening across genome-wide, 47 tomato aquaporins identified nine top-ranked complexes spanning multiple aquaporin subfamilies (PIP1;7, PIP2;1, TIP1;1, TIP2;2, NIP4;1, SIP2;1, XIP1;1, XIP1;3, and XIP1;5). This suggests that harpins may exhibit broader receptor compatibility. Subsequent MD simulation analyses provided critical dynamic and energetic insights. Among the selected complexes, PIP-containing systems (particularly PIP2;1 and PIP1;7) consistently demonstrated high structural stability, compactness, and well-defined free energy minima, supporting previous literature that PIPs are primary and favorable interaction partners. Taken together, the results provide strong computational evidence that: (i) PIP isoforms remain the most probable primary interactor (receptors) of harpin in tomato, (ii) the harpin elicitor is capable of interacting with member from multiple aquaporin subfamilies beyond PIPs, and (iii) specific non-PIP members (particularly TIP and NIP representatives) may act as alternative or auxiliary interaction partners. From a biological perspective, this broad interaction profile of harpin elicitor may reflect a more complex recognition mechanism in plant–pathogen interactions, where different harpin elicitor proteins target multiple membrane channels to drive and modulate diverse cellular responses specific to different non-host plants.

## Abbreviations

AQPs: Aquaporins
FEL: Free energy landscape
MDS: Molecular dynamics simulation
NIP: Nod like intrinsic protein
PCA: Principal component analysis
PIP: Plasma membrane intrinsic protein
PDB: Protein data bank
R_g_: Radius of gyration
RMSD: Root mean square deviations
RMSF: Root mean square fluctuations
SASA: Solvent accessibility surface area
SIP: Small intrinsic protein
TIP: Tonoplast intrinsic protein
T3SS: Type III secretion system
XIP: X- intrinsic protein

## Acknowledgements

We acknowledge the support received from the SiBIOLEAD Team, Little Rock, Arkansas, USA for the MD simulations study as a paid service. We acknowledge the Department of Biotechnology (DBT), GoI, for providing the DBT-JRF fellowship to KL, and stipends to M.Sc. Biotechnology students (TS, SA, GK, AM).

## Author contributions

Conceptualization: DD; Methodology: KL, TS, SA, GK, AM, DD; Formal analysis: KL, DD; Funding acquisition: DD; Project administration: DD; Resources: DD; Supervision: DD; Validation: KL, DD; Visualization: KL, Writing-original draft: KL; Writing-review & editing: KL, DD.

## Data availability

The key data findings generated in this study are available in this article. Any other raw data can be accessed from the corresponding author upon request.

## References

Afzal, Z., Howton, T., Sun, Y., Mukhtar, M., 2016. The roles of aquaporins in plant stress responses. J. Dev. Biol. 4, 9. 10.3390/jdb4010009

Anderson, J.P., Gleason, C.A., Foley, R.C., Thrall, P.H., Burdon, J.B., Singh, K.B., 2010. Plants versus pathogens: an evolutionary arms race. Funct. Plant Biol. 37(6), 499–512. 10.1071/FP09304

Arlat, M., Gijsegem, F.V., Huet, J.C., Pernollet, J.C., Boucher, C.A., 1994. PopA1, a protein which induces a hypersensitivityLlike response on specific *Petunia* genotypes, is secreted via the Hrp pathway of *Pseudomonas solanacearum*. EMBO J. 13(3), 543–553. 10.1002/j.1460-2075.1994.tb06292.x

Bailey, T.L., Johnson, J., Grant, C.E., Noble, W.S., 2015. The MEME suite. Nucleic Acids Res. 43, W39–W49. 10.1093/nar/gkv416

BalintLKurti, P. 2019. The plant hypersensitive response: concepts, control and consequences. Mol. Plant Pathol. 20, 1163–1178. 10.1111/mpp.12821

Barny, M-A., 1995. *Erwinia amylovora* hrpN mutants, blocked in harpin synthesis, express a reduced virulence on host plants and elicit variable hypersensitive reactions on tobacco. Eur. J. Plant Pathol. 101, 333–340. 10.1007/BF01874789

Barrieu, F., Chaumont, F., Chrispeels, M. J., 1998. High expression of the tonoplast aquaporin ZmTIP1 in epidermal and conducting tissues of maize. Plant Physiol. 117(4), 1153–1163. 10.1104/pp.117.4.1153

Bowie, J.U., Lüthy, R., Eisenberg, D., 1991. A method to identify protein sequences that fold into a known three-dimensional structure. Science. 253, 164–170. 10.1126/science.1853201

Büttner, D., He, S.Y., 2009. Type III protein secretion in plant pathogenic bacteria. Plant Physiol. 150, 1656–1664. 10.1104/pp.109.139089

Chen, L., Qian, J., Qu, S., Long, J., Yin, Q., Zhang, C., et al., 2008. Identification of specific fragments of HpaG_Xooc_, a harpin from *Xanthomonas oryzae* pv. *oryzicola*, that induce disease resistance and enhance growth in plants. Phytopathology. 98, 781–791. 10.1094/PHYTO-98-7-0781

Choi, M-S., Kim, W., Lee, C., Oh, C-S., 2013. Harpins, multifunctional proteins secreted by gram-negative plant-pathogenic bacteria. Mol. Plant-Microbe Interact. 26, 1115–1122. 10.1094/MPMI-02-13-0050-CR

Colovos, C., Yeates, T.O., 1993. Verification of protein structures: patterns of nonbonded atomic interactions. Protein Sci. 2, 1511–1519. 10.1002/pro.5560020916

Gasteiger, E., Hoogland, C., Gattiker, A., Duvaud, S., Wilkins, M.R., Appel, R.D., et al., 2005. Protein identification and analysis tools on the ExPASy server. In: Walker JM (ed), The proteomics protocols handbook. Humana Press, Totowa, NJ, pp 571–607. 10.1385/1-59259-890-0:571

Gaudriault, S., Brisset, M-N., Barny, M-A., 1998. HrpW of *Erwinia amylovora*, a new Hrp-secreted protein. FEBS Lett. 428, 224–228. 10.1016/S0014-5793(98)00534-1

He, S. Y., Huang, H. C., and Collmer, A., 1993. *Pseudomonas syringae* pv. *syringae* harpin*_Pss_*: a protein that is secreted via the Hrp pathway and elicits the hypersensitive response in plants. Cell. 73(7), 1255–1266. 10.1016/0092-8674(93)90354-S

Hukin, D., Doering-Saad, C., Thomas, C. and Pritchard, J., 2002. Sensitivity of cell hydraulic conductivity to mercury is coincident with symplasmic isolation and expression of plasmalemma aquaporin genes in growing maize roots. Planta. 215(6), 1047–1056.

Jia, L., Zhu, L., 2025. The bacterial type III secretion system as a broadly applied protein delivery tool in biological sciences. Microorganisms. 13(1), 75. 10.3390/microorganisms13010075

Jones, G., Jindal, A., Ghani, U., Kotelnikov, S., Egbert, M., Hashemi, N., et al., 2022. Elucidation of protein function using computational docking and hotspot analysis by ClusPro and FTMap. Acta Crystallogr. D Struct. Biol. 78, 690–697. 10.1107/S2059798322002741

Kaur, S., Samota, M.K., Choudhary, M., Choudhary, M., Pandey, A.K., Sharma, A., et al., 2022. How do plants defend themselves against pathogens-Biochemical mechanisms and genetic interventions. Physiol. Mol. Biol. Plants. 28, 485–504. 10.1007/s12298-022-01146-y

Kim, J.F., Beer, S.V., 1998. HrpW of *Erwinia amylovora*, a new harpin that contains a domain homologous to pectate lyases of a distinct class. J. Bacteriol. 180, 5203–5210. 10.1128/JB.180.19.5203-5210.1998

Kolodny, R., Kosloff, M., 2013. From protein structure to function via computational tools and approaches. Israel J. Chem. 53, 147–156. 10.1002/ijch.201200078

Kourghi, M., Pei, J.V., De Ieso, M.L., Nourmohammadi, S., Chow, P.H., Yool, A.J., 2018. Fundamental structural and functional properties of aquaporin ion channels found across the kingdoms of life. Clin. Exp. Pharmacol Physiol. 45(4), 401–409. 10.1111/1440-1681.12900

Lal, K., Joshi, A., Saini, V., Mohammed, M., Sarma, P.V.S.R.N., Dey, D., 2026. Molecular identification and functional analysis of HrpZ2, a new member of the harpin superfamily from *Pseudomonas syringae*, inducing hypersensitive response in tobacco. Front. Plant Sci. 16, 1665817. 10.3389/fpls.2025.1665817

Lal, K., Singh, V.K., Dey, D., 2023. Study of molecular interaction between *Arabidopsis* aquaporin and *Pseudomonas syringae* pv. *syringae* harpin (HrpZ*_Pss_*) through molecular docking tools. Int. J. Plant Env. 9, 343–349. 10.18811/ijpen.v9i04.06

Laskowski, R.A., Rullmann, J.A.C., MacArthur, M.W., Kaptein, R., Thornton, J.M., 1996. AQUA and PROCHECK-NMR: programs for checking the quality of protein structures solved by NMR. J. Biomol. NMR. 8. 10.1007/BF00228148

Letunic, I., Bork, P., 2024. Interactive tree of life (iTOL) v6: recent updates to the phylogenetic tree display and annotation tool. Nucleic Acids Res. 52, W78–W82. 10.1093/nar/gkae268

Li, J., Liu, H., Cao, J., Chen, L-F., Gu, C., Allen, C., et al., 2010. PopW of *Ralstonia solanacearum*, a new twoLJdomain harpin targeting the plant cell wall. Mol. Plant Pathol. 11, 371–381. 10.1111/j.1364-3703.2010.00610.x

Li, L., Wang, H., Gago, J., Cui, H., Qian, Z., Kodama, N., et al., 2015. Harpin Hpa1 interacts with aquaporin PIP1;4 to promote the substrate transport and photosynthesis in *Arabidopsis*. Sci. Rep. 5,17207. 10.1038/srep17207

Li, P., Zhang, L., Mo, X., Ji, H., Bian, H., Hu, Y., et al., 2019. Rice aquaporin PIP1;3 and harpin Hpa1 of bacterial blight pathogen cooperate in a type III effector translocation. J. Exp. Bot. 70, 3057–3073. 10.1093/jxb/erz130

Liu, C., Fukumoto, T., Matsumoto, T., Gena, P., Frascaria, D., Kaneko, T., et al., 2013. Aquaporin OsPIP1; 1 promotes rice salt resistance and seed germination. Plant Physiol. Biochem. 63, 151–158. 10.1016/j.plaphy.2012.11.018

Liu, Y., Zhou, X., Liu, W., Xiong, X., Lv, C., Zhou, X., Miao, W., 2018. Functional regions of HpaXm as elicitors with specific heat tolerance induce the hypersensitive response or plant growth promotion in nonhost plants. PLoS One. 13(1), e0188788. 10.1371/journal.pone.0188788

Patoliya, J., Thaker, K., Rabadiya, K., Patel, D., Jain, N. K., Joshi, R., 2024. Uncovering the Interaction Interface Between Harpin (Hpa1) and Rice Aquaporin (OsPIP1;3) Through Protein-Protein Docking: An In Silico Approach. Mol Biotechnol. 66(4), 756–768. 10.1007/s12033-023-00690-6

Sukhwal, A., Sowdhamini, R., 2015. PPCheck: A webserver for the quantitative analysis of protein-protein interfaces and prediction of residue hotspots. Bioinform. Biol. Insights. 9, BBI.S25928. 10.4137/BBI.S25928

Tian, S., Wang, X., Li, P., Wang, H., Ji, H., Xie, J., et al., 2016. Plant aquaporin AtPIP1; 4 links apoplastic H2O2 induction to disease immunity pathways. Plant Physiol. 171(3), 635–1650. 10.1104/pp.15.01237

Verkman, A.S., 2008. Mammalian aquaporins: diverse physiological roles and potential clinical significance. Expert Rev. Mol. Med. 10, e13. 10.1017/S1462399408000690

Wang, E., Fu, W., Jiang, D., Sun, H., Wang, J., Zhang, X., et al., 2021. VAD-MM/GBSA: a variable atomic dielectric MM/GBSA model for improved accuracy in protein–ligand binding free energy calculations. J. Chem. Inf. Model. 61, 2844–2856. 10.1021/acs.jcim.1c00091

Wang, X., Li, M., Zhang, J., Zhang, Y., Zhang, G., Wang, J., 2007. Identification of a key functional region in harpins from *Xanthomonas* that suppresses protein aggregation and mediates harpin expression in *E. coli*. Mol. Biol. Rep. 34, 189–198. 10.1007/s11033-006-9034-6

Wang, Y., Zhao, Z., Liu, F., Sun, L., Hao, F., 2020. Versatile roles of aquaporins in plant growth and development. Int. J. Mol. Sci. 21, 9485. 10.3390/ijms21249485

Wei, Z-M., Laby, R.J., Zumoff, C.H., Bauer, D.W., He, S.Y., et al., 1992. Harpin, elicitor of the hypersensitive response produced by the plant pathogen *Erwinia amylovora*. Science. 257, 85–88. 10.1126/science.1621099

Xu, F., Wang, K., Yuan, W., Xu, W., Liu, S., Kronzucker, H.J., et al., 2019. Overexpression of rice aquaporin OsPIP1; 2 improves yield by enhancing mesophyll CO_2_ conductance and phloem sucrose transport. J. Exp. Bot. 70(2), 671–681. 10.1093/jxb/ery386

Zhang, J., Qian, J., 2024. Advances in computational intelligence-based methods of structure and function prediction of proteins. Biomolecules. 14(9), 1083. 10.3390/biom14091083

Zheng, W., Wuyun, Q., Li, Y., Liu, Q., Zhou, X., Peng, C., et al., 2026. Deep-learning-based single-domain and multidomain protein structure prediction with D-I-TASSER. Nat. Biotechnol. 44, 641–653. 10.1038/s41587-025-02654-4

